# High concordance between genetic effects on mRNA and protein abundance

**DOI:** 10.64898/2026.05.26.727960

**Authors:** Krisna Van Dyke, Matthew Feraru, Frank W. Albert

**Affiliations:** Department of Genetics, Cell Biology, & Development, University of Minnesota, Minneapolis 55455, MN, USA; Department of Medical Oncology, Dana-Farber Cancer Institute, Boston, 02115, MA, USA; Apple Inc., Cupertino 95014, CA, USA

## Abstract

Genetic influences on gene expression are an important source of variation in complex traits. Conflicting results have been reported about the concordance of genetic effects on mRNA abundance vs. protein levels, ranging from high agreement to a predominance of effects that are specific to mRNA or protein. Here, we integrated 13 published datasets of genetic variation in mRNA or protein collected in the same cross of two strains of the yeast *Saccharomyces cerevisiae*. These highly replicated data allowed us to gauge the overall agreement between the genetics of mRNA and protein and search for individual loci whose effects on these two gene products are reproducibly different. Overall, genetic effects were highly correlated across all datasets. mRNA and protein showed similar genetic architectures. Pairwise agreement between loci from mRNA datasets and loci from protein datasets was indistinguishable from agreement between loci from datasets of the same gene product. *Trans*-acting hotspots with effects on numerous genes affected mRNA and protein similarly. There were no hotspots that exclusively affected mRNA or protein across datasets. A small number of loci did show reproducibly different effects on mRNA or protein of individual genes. Collectively, these results show that, with a few notable exceptions, genetic effects on mRNA and protein are largely concordant.

## Introduction

Genetic effects on gene expression are an important influence on complex traits [1]. To map this regulatory genetic variation, numerous studies have measured gene expression in sets of genetically different individuals to identify quantitative trait loci (QTLs) that harbor DNA variants that affect the expression of a given gene [2].

Due to practical reasons of low cost, high throughput, and the ability to quantify the expression of most genes in the genome, most studies of regulatory variation use mRNA abundance as a measure of gene expression. So far, this work has culminated in large, comprehensive catalogs of “expression QTLs” (eQTLs) in a range of species [3–9].

This rich body of literature has revealed key insights into the genetic architecture of variation in gene expression. Regulatory variation is pervasive, such that the expression of nearly all genes is affected by at least one eQTL [3,5,6]. Local eQTLs arise from DNA variants located close to the gene they affect, where they primarily (but not always [10,11]) alter *cis*-regulatory elements to change transcription or mRNA stability [11,12]. By contrast, distant eQTLs arise from variants that alter the activity or abundance of *trans*-acting regulatory genes, which in turn affect the expression of other genes that can be located anywhere in the genome [13,14]. Individual local eQTLs tend to have larger effects than individual *trans*-eQTLs, making them easier to discover in studies with limited statistical power. However, most genes are affected by multiple *trans*-eQTLs [5,15], which collectively contribute more gene expression heritability than do local eQTLs [5,16,17]. *Trans*-eQTLs for multiple genes tend to co-occur at certain “hotspot” locations [5,13,14,18] caused by variation in genes with varied functions ranging from transcription factors to metabolic regulators and enzymes [19–22]. Genetic influences on mRNA abundance can vary among contexts such as tissues, cell types, developmental stage, and cellular environments [3,23–26].

Proteins are the functional product of many genes. Following pioneering work exploring the genetic basis of variation in protein levels [18,27–29], extensive sets of “protein QTLs” (pQTLs) now exist, e.g. from human plasma [30–33], brain [34], and cell lines [35], a diverse panel of natural yeast isolates [36], highly powered crosses between pairs of yeast strains [21,37], and diverse mouse intercross panels [28,38].

A key question motivating pQTL mapping is the similarity of genetic effects on mRNA and protein. Does a typical eQTL also result in a pQTL for the same gene because the altered mRNA abundance is carried forward to a difference in protein level? Or do buffering mechanisms predominate, such that protein levels are unaffected by genetic influences on mRNA abundance [18,28,34,35,39,40]? How common are pQTLs that arise from protein-specific mechanisms such as mRNA translation [35,41] and protein degradation [42–44] that are independent of the gene’s mRNA abundance?

The results so far have been inconclusive and conflicting. On the one hand, pQTLs recapitulate general patterns seen in eQTLs. Similar numbers of genes show local eQTLs and pQTLs [36,37,45], and most local pQTLs appear to arise from differences in mRNA [38]. There are similar numbers of eQTLs and pQTLs per gene [5,21], with numerous *trans*-pQTLs that are individually weaker [36,37,46] but collectively account for more variation in protein levels than local pQTLs [21,46]. *Trans*-pQTL hotspots tend to occur at the same locations as *trans*-eQTL hotspots [21,37,45,47]. Increasing the statistical power of pQTL mapping can result in the discovery of many pQTLs at eQTLs that had previously appeared to be mRNA-specific [37]. A comparison between well-powered eQTL and pQTL datasets [5], as well as a study mapping eQTLs and pQTLs in the same yeast growth cultures [45] found strong correlations between the effect sizes of eQTLs and pQTLs affecting the same gene.

On the other hand, considerable discrepancies between the genetics of the two layers of gene expression have been observed. Local pQTLs can arise from missense variants in the coding region of a gene that destabilize the protein but do not strongly affect the gene’s mRNA [21]. Some studies reported fewer and weaker local pQTLs than local eQTLs [48,49], perhaps due to pervasive buffering of eQTLs at the protein level [34,35]. Low or even negative correlations between mRNA and protein levels may arise from distinct sets of local eQTLs and local pQTLs [50].

Even more severe discrepancies have been reported between *trans*-eQTLs and *trans*-pQTLs. In recombinant mouse panels, *trans*-eQTLs and *trans*-pQTLs were found to have very little overlap, and *trans*-pQTL were deemed to arise largely independently of mRNA variation [28,38]. Similarly, a yeast genome-wide association study found almost no overlap between *trans*-pQTLs and *trans*-eQTLs [36]. In a yeast cross, a powerful bulk-segregant approach that quantified mRNA and protein of ten genes in the same cells found a majority of *trans*-QTLs to be specific for protein or mRNA [51]. *Trans*-QTL hotspots that are entirely or largely specific to mRNA or protein have been reported [18,27,37,45]. Even when hotspots do affect both mRNA and protein, their effects on the two products of the same genes can be quite different [5,27,45,49]. Mechanisms causing these discrepancies may involve *trans*-acting influences on the protein degradation machinery [42,43], kinases [51], indirect mechanisms that shape amino acid metabolism, ribosome production, and translation [45,49]; or degradation of protein complex members due to stoichiometric imbalance in protein-protein interactions [38,46].

These conflicting results on the similarity between genetic influences on mRNA versus protein pose major questions. If mRNA abundance and eQTLs are typically not informative for proteins, why should they be studied, given that proteins are the functional gene product? Conversely, if protein levels and pQTLs are well-approximated by mRNA, why invest in costly and challenging quantification of proteins in large samples?

A unique collection of datasets in the yeast *Saccharomyces cerevisiae* promises to shed light on this issue. Since 2002, a cross between the laboratory strain BY and the vineyard strain RM has been used in at least 13 studies mapping genetic influences on mRNA abundance, protein levels, or both (Supplementary Table S1). In this cross, meiotic recombination creates haploid segregants that each carry a different, random combination of BY and RM alleles. Technologies used to quantify gene expression in these segregants have ranged from microarrays and RNA-seq for mRNA to various mass-spectrometry strategies and green fluorescent protein (GFP) tags for proteins. Mapping strategies have included “individual-segregant” panels [52], in which segregants are phenotyped and genotyped separately, as well as bulk-segregant approaches using large pools of recombinant cells [37,47,51]. Crucially, all of these studies were conducted with the same two parent strains and sometimes even the same set of segregants.

In spite of this extensive body of work, there is no consensus on the degree of discrepancy between the genetics of mRNA vs protein in this cross. Partly, this is because each new dataset was compared to just one other dataset, either from a previous publication or from the same project. Each eQTL and pQTL study has idiosyncrasies, including environmental variation that exists even when experimental conditions are carefully controlled (for example due to different media batches), the exact identity of the analyzed individuals, as well as different measurement technologies and QTL mapping algorithms. Together with stochastic biological variation, measurement noise, and incomplete statistical power, these idiosyncrasies may skew comparisons between any two datasets. Integrative analyses of multiple datasets could provide more rigorous assessments. However, no study so far has examined the collective corpus of information available in the BY/RM cross.

Here, we integrated data from nearly all BY/RM eQTL and/or pQTL studies to date. We asked two questions: how similar are genetic effects on mRNA and protein? And where are loci with the most convincing evidence for mRNA or protein-specific effects? We found high overall concordance among datasets, similar genetic architectures for mRNA and protein, and only a few loci with significantly different effects on the two gene products. A parsimonious interpretation of these results is that the effects of regulatory variation on mRNA and protein are typically the same, with small numbers of noteworthy exceptions.

## Results

### Dataset overview

We analyzed 13 BY/RM datasets (7 mRNA and 6 protein) from 10 publications (Table 1 & Supplementary Table S1). In all datasets, gene expression was measured during exponential growth. Most datasets (n = 9) used synthetic complete medium with glucose (SC), while the remaining four used yeast nitrogen base (YNB) medium with either glucose (n = 3) or ethanol (n = 1) as the carbon source.

**Table 1.**
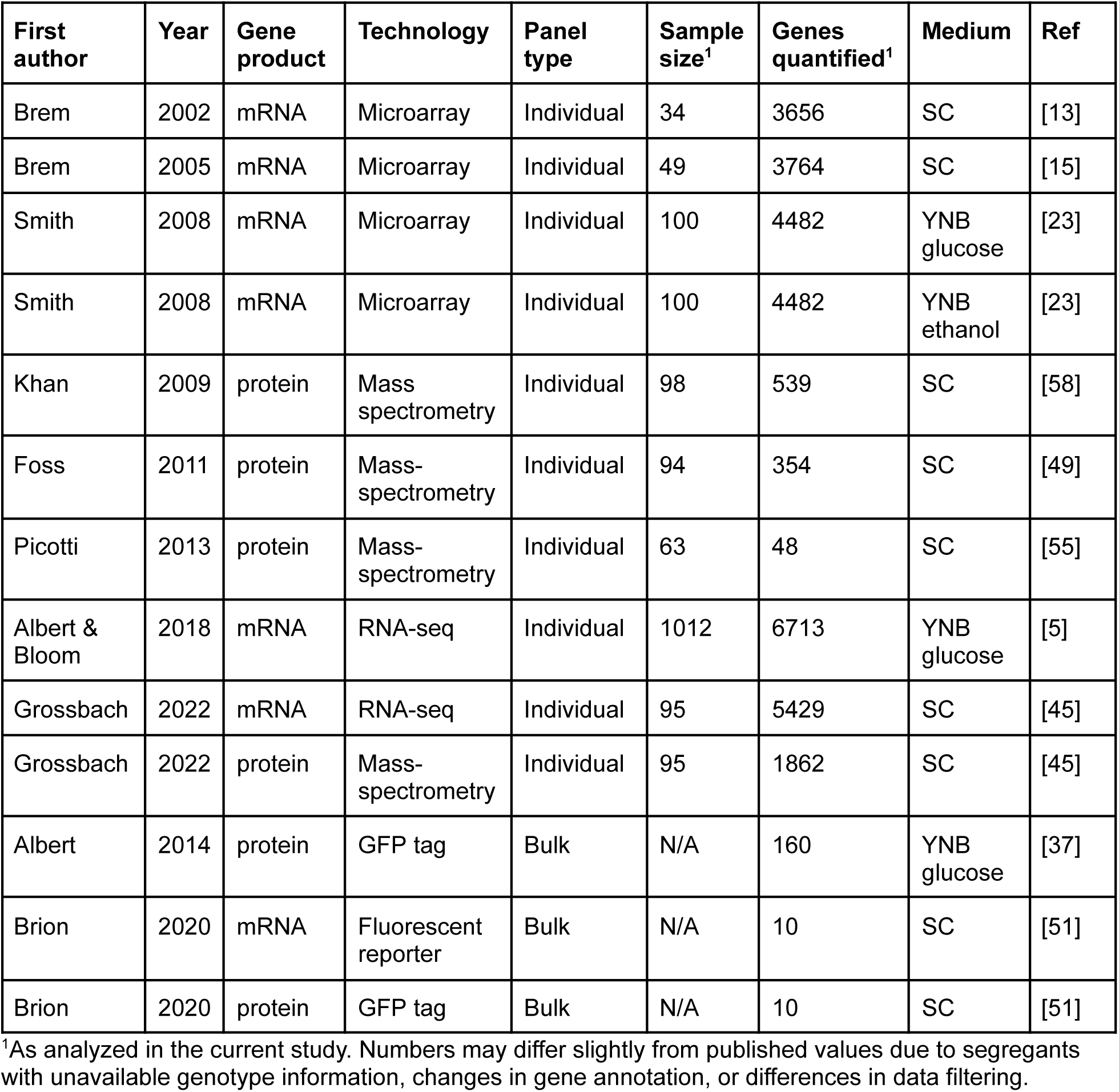
Datasets analyzed in this paper.

The individual-segregant design was used in 10 datasets (6 mRNA and 4 protein, 8 publications). Across these, mRNA measurements were available for 3,656 to 6,713 genes, protein levels were available for 48 to 1,862 genes, and sample sizes ranged from 34 to 1,012 segregants. All individual-segregant studies except for Albert & Bloom et al., 2018, which used a newer segregant set [53], used the same panel of BY / RM segregants. This panel, published in Brem et al., 2005 [15] as an expansion of an earlier set [13], was later re-genotyped by integration of RNA-seq data by Grossbach et al. [45], yielding more complete coverage of the genome. In our analyses, we used these improved genotypes at 3,593 unique markers.

We also incorporated three sets of QTLs that had been mapped using bulk-segregant analysis [37,51]. In this approach, a gene of interest is tagged with a fluorescent reporter of mRNA and / or protein abundance in one or both parent strains. The parents are crossed to generate pools of recombined cells that are then sorted into populations with high or low expression of the given gene. Whole-genome sequencing identifies QTLs affecting the abundance of the tagged gene product as allele frequency differences between these two populations. This design provides high statistical power but is not compatible with individual-segregant mapping pipelines. Therefore, we used the published pQTLs for 160 proteins from Albert et al., 2014 [37] and eQTLs and pQTLs for 10 genes from Brion et al., 2020 [51] wherever possible.

### Multivariate modeling reveals broad sharing of genetic effects across datasets

We first sought to gauge the similarity of genetic effects across studies globally, without considering individual QTLs. To do so, we analyzed the individual-segregant datasets using multivariate adaptive shrinkage (mashr [54]). Mashr models relationships among datasets by learning correlation patterns among the genetic effects in each dataset. We trained mashr on linkage disequilibrium (LD)-pruned gene-marker pairs from 385 genes represented in at least two protein and two mRNA datasets. A mixture of gene-marker pairs with strong genetic signals and those from the genomic background were used to stabilize covariance learning (Methods). We then visualized the degree of effect sharing via a correlation matrix and analyzed patterns of effect sharing using principal component analysis (Figure 1).

**Figure 1.**
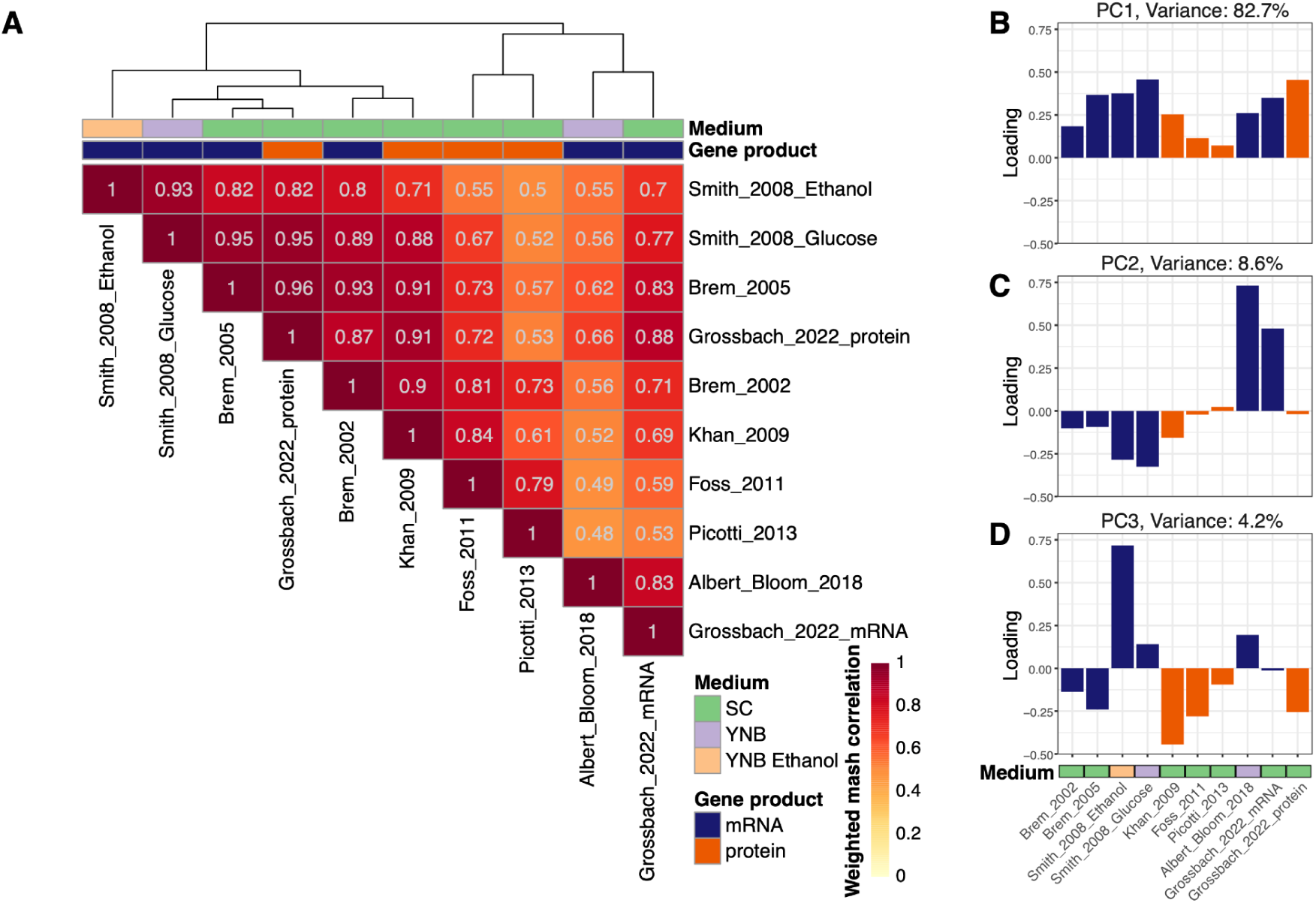
Global sharing of effects across datasets. A. Correlation matrix of effect sharing among datasets. The matrix is clustered based on similarity. B. Principal components (PCs) of global effect sharing. Loadings of each dataset onto the first three PCs are shown.

Similarity of genetic effects across datasets was overall high, with a median correlation of 0.73 and positive correlations among all datasets (Figure 1A). This pattern was predominant, accounting for 83% of the variance (Figure 1B). Additional features were apparent. The two RNA-sequencing datasets (Albert & Bloom et al. and Grossbach et al.) [5,45] were highly correlated with each other and stood out from the other datasets (Figure 1C). Datasets collected in SC medium were differentiated from those collected in YNB (Figure 1D). By contrast, mRNA and protein datasets were not well separated by any principal component (Supplementary Figure S1; also note that mRNA and protein datasets did not cluster in the correlation matrix in Figure 1A). These results suggest that genetic effects on gene expression are generally shared across these 10 datasets. In particular, genetic effects on mRNA and protein appeared to be quite similar.

### Numerous *trans*-acting loci shape mRNA and protein levels

To dissect the basis of these global patterns, we mapped QTLs in all individual-segregant panels. We used the mapping pipeline from Albert & Bloom et al. [5], which employs chromosome-specific forward scans that control for genetic variation on other chromosomes while using permutations to estimate the false-discovery rate (FDR).

Across the individual-segregant studies, we identified a median of 1,018 QTLs per dataset (range: 16 pQTLs for 13 out of 48 proteins measured in Picotti et al. [55] to 36,723 eQTLs for 5,644 out of 6,713 mRNAs quantified in Albert & Bloom et al. [5]) at an FDR of 5% (Supplementary Table S2; QTLs are also available for interactive exploration in the UCSC genome browser at https://genome.ucsc.edu/s/kjv/BYxRM_all_QTL_tracks). The QTLs showed good agreement with available published QTL lists (Supplementary Table S3). Thus, our re-mapped QTLs recapitulate published results via an identical mapping pipeline, eliminating variation that may arise from different QTL mapping algorithms.

Larger sample size resulted in more QTL discoveries, as expected given higher statistical power (Figure 2A, analysis of variance (ANOVA) of a negative binomial model of QTL counts, p = 1.7e-10 using all individual-segregant datasets; p = 7.9e-7 without the large Albert & Bloom dataset). eQTLs and pQTLs were discovered at similar rates as a function of sample size (interaction between gene product and sample size: p ≥ 0.29). We classified QTLs as “local” if they included their target gene (Methods; this terminology allows for the existence of local *trans* effects [5,10] rather than assume that all local QTLs act in *cis*) and as *trans* otherwise. In all datasets, there were more *trans*-QTLs than local QTLs (Figure 2B). *Trans*-QTLs were discovered at a higher rate than local QTLs as a function of sample size (p = 0.0002); this effect was not detectable when the large Albert & Bloom eQTL study was excluded (p = 0.17). The rates of local vs *trans*-QTL discovery as a function of sample size were not different between mRNA and protein (p ≤ 0.60). Individual local QTLs had stronger effects than individual *trans*-QTLs, and larger sample size resulted in the detection of QTLs of smaller effect, as expected [56] (Supplementary Figure S2). Published QTLs from bulk-segregant studies reflected these patterns (Supplementary Text 1). Collectively, these analyses agree with recent work [5,21,37] showing that mRNA and protein levels have similar genetic complexity, with prominent roles for *trans*-acting variation in shaping both mRNA and protein abundance.

**Figure 2.**
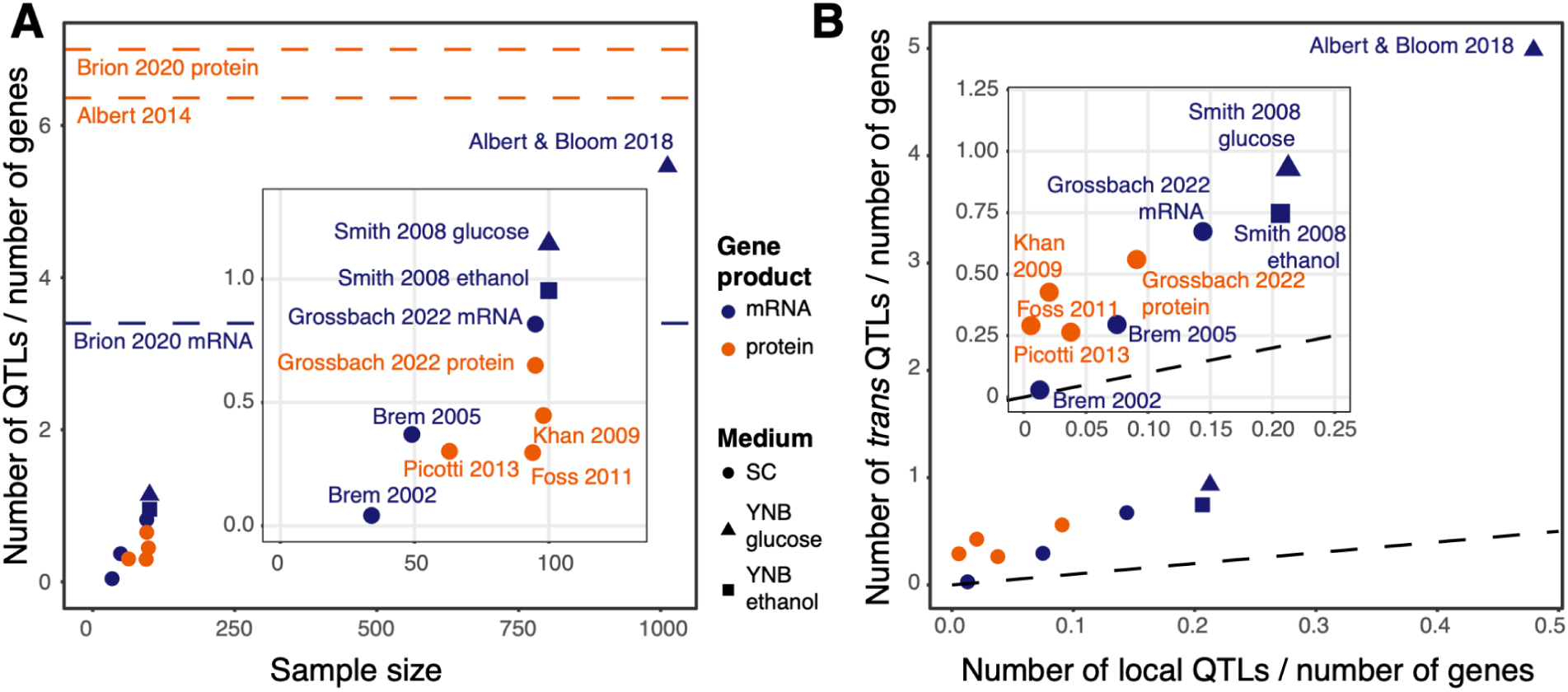
Patterns of QTL discovery across datasets. A. QTL discovery as a function of sample size. The y-axis shows the total number of QTLs divided by the number of analyzed genes in each study. The inset excludes the large Albert & Bloom et al. eQTL study. Dashed horizontal lines show the number of QTLs per gene discovered in bulk-segregant datasets (Supplementary Text 1). B. Numbers of local versus *trans*-QTLs per gene. The black dashed line indicates an equal number of local and *trans*-QTLs per gene. Note that all datasets are above this line. Bulk segregant data for 41 proteins follows the same trend (Supplementary Text 1).

### QTL agreement between mRNA and protein datasets is similar to that among mRNA and among protein datasets

Next, we quantified how well QTLs between different datasets agree with each other. Our analyses were motivated by the wide range of such QTL agreements reported in the literature. For example, 60% of the eQTLs found by Smith & Kruglyak, 2008 in glucose [23] were later reported to correspond to a pQTL in the Albert et al., 2014 bulk-segregant study [37]. Likewise, around 50% agreement between strong QTLs was reported between eQTLs from Albert & Bloom et al., 2018 [5] and pQTLs from Albert et al., 2014 [37]. On the other hand, pQTL / eQTL agreement was reported to range from only 20% down to no more than random for certain gene groups in Foss et al., 2011 [49], and was reported to be only 3% in Teyssonnière et al., 2024 [36]. These published comparisons all contrasted one newly collected dataset to one other dataset, each providing only a single observation of QTL set agreement.

To quantify QTL agreement more fully, we compared QTLs between all pairs of datasets. For each pair, we calculated the fraction of QTLs identified in one dataset (the “query” set) that matched a QTL in the “target” dataset. Target QTLs were deemed to match a query if they were genome-wide significant in the target study, had confidence intervals overlapping the query QTL, and had the same direction of effect. The average agreement between mRNA queries and protein targets was 20.8%, with a wide range (Figure 3). The average agreement between protein and mRNA datasets was 15.0%. While these two values may at first suggest that more eQTLs tend to correspond to a pQTL than *vice versa*, the distributions of agreements across all dataset pairs were not significantly different (binomial mixed model: p = 0.57; Supplementary Table S4).

**Figure 3:**
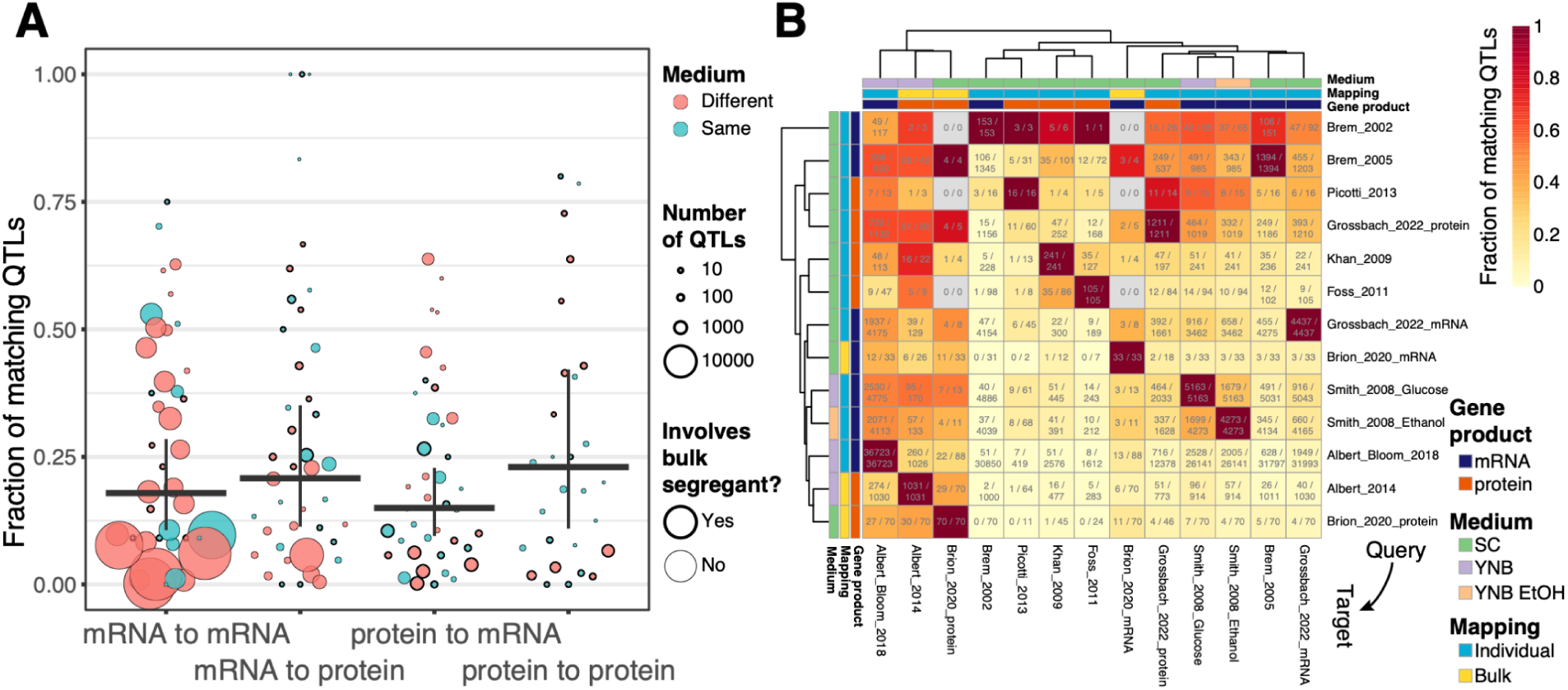
Pairwise comparisons between datasets. A. Fraction of QTLs in a query set that matches a QTL in a given target set. Circles are sized by the number of QTLs in the given comparison. Thick horizontal lines show the mean for each distribution, weighted by dataset sizes as computed using linear models that control for dataset identity (Methods). Vertical lines are standard errors of these means, computed from the same models. B. Agreements for each pairwise comparison, arranged as a heatmap. Numbers in cells are the number of query QTLs that match a QTL in the target set, along with the number of all query QTLs. Gene product, mapping method, and growth medium are indicated. Rows show query datasets, and columns show target datasets. Rows and columns are clustered based on similarity.

To provide context to these observations, we examined agreement between datasets of the *same* gene product. If a large fraction of QTLs affect only the mRNA or only the protein of a given gene (or affect both gene products, but in different directions), we would expect to see less agreement between datasets of different gene products (i.e., eQTLs versus pQTLs) than between pairs of datasets of the same gene product (i.e. two eQTL datasets, or two pQTL datasets). No such difference was seen.

Specifically, the average agreement between two eQTL datasets was 17.9%, and the average agreement between two pQTL datasets was 23.0%. Neither agreement was significantly different from those between datasets of different gene products (p ≥ 0.41; Supplementary Table S4). These results were robust to restricting the analyses to the strongest QTLs, to datasets sharing the same medium, to individual-segregant panels, and to local QTLs. Similar results were obtained using the π1 statistic for QTL replication [57], as well as using correlations of QTL effect sizes as a measure of agreement (Supplementary Text 2, Supplementary Figure S3; Supplementary Table S4).

These results show that differences between sets of eQTLs and pQTLs do not exceed a random expectation formed by comparing eQTLs to eQTLs or pQTLs to pQTLs. Differences among QTLs for the same gene product likely reflect incomplete statistical power, different measurement techniques, slight differences in environment, and random biological fluctuations. All these factors also affect comparisons of different gene products. True differences in genetic effects on mRNA vs. protein would be expected to create excess discrepancy and less agreement. This was not observed, arguing against the existence of large numbers of genetic effects that specifically influence mRNA or protein.

### *Trans*-acting hotspots are not specific to mRNA or protein

Given their prominence in regulatory variation, we examined evidence for mRNA or protein specificity at *trans*-QTLs. *Trans*-QTLs tend to reside at hotspot locations that affect the expression of multiple genes (Figure 4). We expressed the strength of a hotspot in a given dataset as the fraction of QTLs in that dataset whose peaks fall into a given genome bin. Figure 5 shows these results using 100 kb-wide bins, while Supplementary Table S5 presents more granular results for bins of 10 kb.

**Figure 4.**
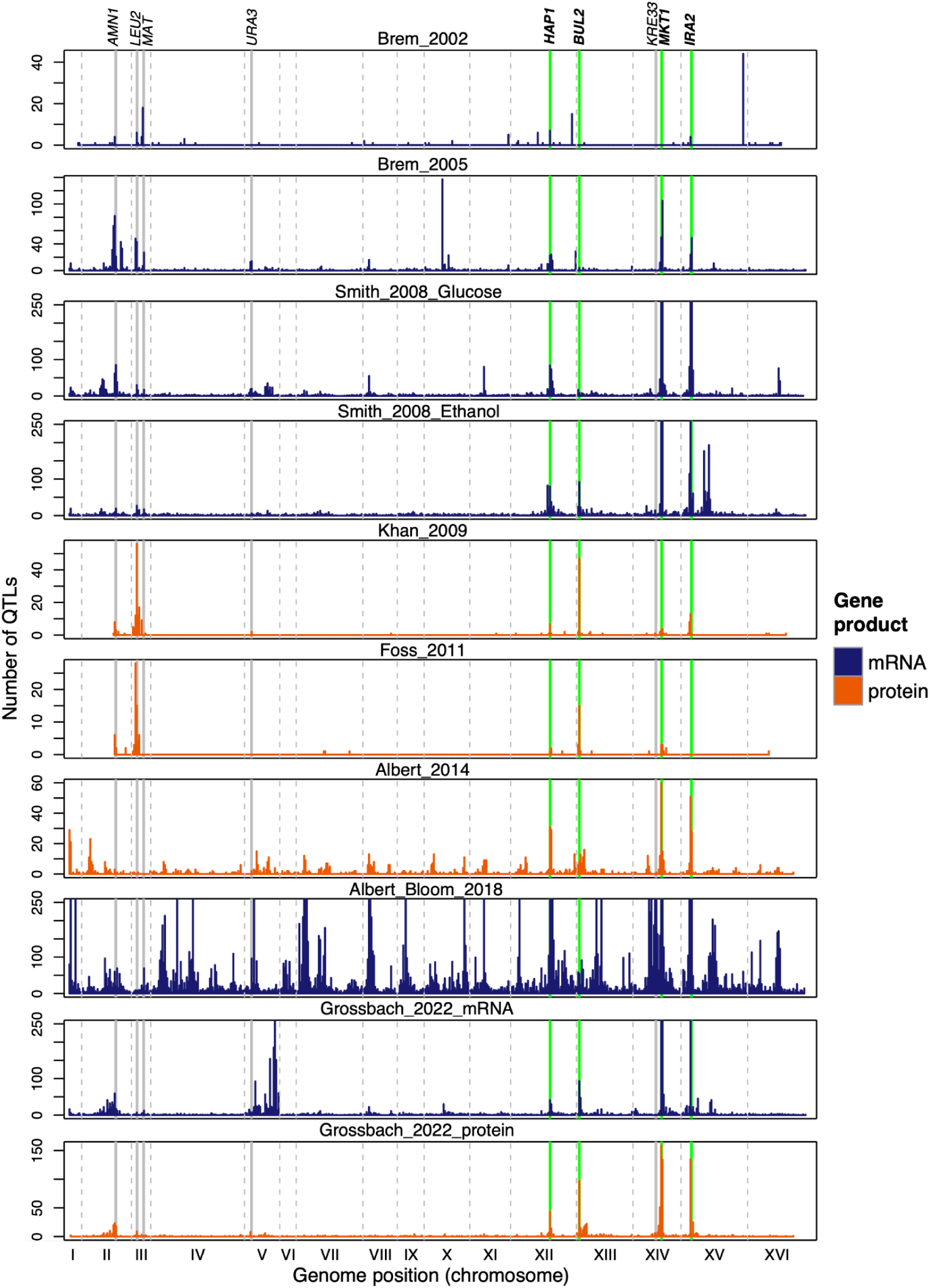
Location of *trans*-QTLs along the genome. Solid green vertical lines and bold gene names: hotspots highlighted in the text whose causal alleles exist in all datasets. Solid grey vertical lines and regular gene names: hotspots whose causal alleles segregate in only some datasets. Dashed grey vertical lines: chromosome boundaries. Shown are datasets with at least analyzed 100 genes. Y-axes are truncated to show a maximum of 250 QTLs.

**Figure 5.**
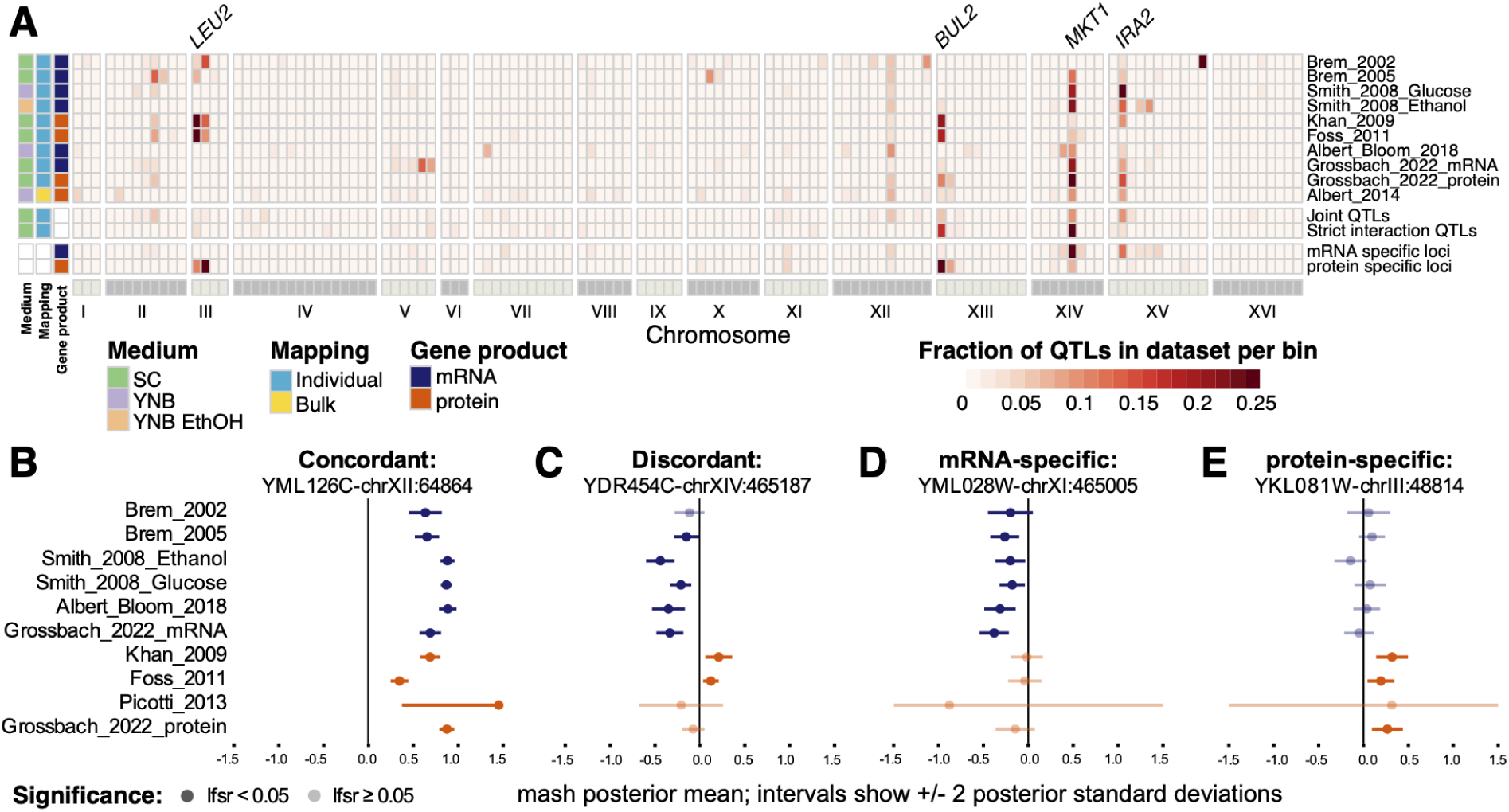
QTLs with distinct effects on mRNA or protein. A. The figure shows the fraction of QTLs in each dataset (rows) whose peak marker falls into a given 100 kb genome bin (columns). QTLs from individual studies with at least 100 genes are shown, along with jointly mapped QTLs and mRNA or protein-specific QTLs. Dataset characteristics (medium, mapping method, and gene product) are indicated on the left. Genes mentioned in the text likely to underlie *trans*-QTL hotspots are indicated. B – E. Examples of jointly-mapped QTLs that affect mRNA and protein of a given gene similarly (B), in opposite directions (C), affect only mRNA (D) or only protein (E). The affected gene and the peak marker of the QTL are shown above each plot. Posterior estimates of effect sizes calculated by mashr are shown.

Several prominent hotspots recurred across studies. These include hotspots at the *HAP1* [13], *MKT1* [58], and *IRA2* genes [20,23] that were present in all or nearly all studies (green vertical lines in Figure 4). They also include hotspots at the mating type locus (*MAT*), *AMN1*, and engineered auxotrophies such as *LEU2* that were present in studies in which the causal allele segregates [13,14,20,58] (Figure 4; the specific BY / RM parents used in some studies did not differ at these causal alleles, and no hotspot is therefore expected at these positions). There was considerable heterogeneity in how many genes these major loci affected. For example, the *IRA2* locus accounted for 28% of the glucose eQTLs from Smith & Kruglyak 2008, [23] but no pQTLs in Foss et al., 2011 [49], with a median across studies of 8%. Thus, *trans*-acting hotspots generally reproduce across studies of both mRNA and protein but with variable strength.

We searched for hotspots with protein- or mRNA-specific effects. We focused on hotspots that appeared to affect only or primarily mRNA or protein at the time of their initial publication. For example, the Grossbach et al., 2022 mRNA dataset has a strong hotspot (15% of QTLs) at 470 - 520 kb on chromosome V. We confirmed that this hotspot is absent in the protein data collected in the same study from the same yeast cultures (there were zero QTLs in this region), raising the possibility that this hotspot could be mRNA-specific. However, this hotspot was also absent or much weaker in all other mRNA datasets (maximum of these studies: 0.09% of the Smith & Kruglyak ethanol eQTLs [23]), including in the Albert & Bloom et al., 2018 mRNA dataset that had ten times larger sample size (0.08% of eQTLs at this locus). Similarly, one of the strongest mRNA hotspots in Brem et al., 2005, with 9% of all QTLs [15] on chromosome X (290 - 300 kb), was absent or much weaker in all other studies, including one using the same segregants and the same medium (Grossbach et al., 2022 mRNA [45]; 0.02%; note that this study did call an mRNA hotspot in this general location, but in our analyses this hotspot was ∼20 kb away (320 kb - 340 kb) from that in Brem et al., 2005). A hotspot on chromosome II (3% of QTLs, 120 - 150 kb) appeared to be largely protein-specific in Albert et al., 2014 [37]. However, our analyses here show that this hotspot was not detected in other protein datasets. Overall, we found no hotspot that affected only mRNA or only protein in all datasets of the given gene product. Thus, the same *trans*-acting hotspots are seen in both mRNA and protein datasets, but with variable strength that may depend on each dataset’s exact environmental conditions. Hotspots whose effects are entirely and reproducibly specific to mRNA or protein were not apparent.

### Joint QTL mapping reveals loci that differentially affect mRNA and protein for individual genes

To examine genetic effects on the expression of individual genes, we implemented mapping pipelines that analyze datasets jointly (Methods). To avoid confounding by environmental variation, we used only the seven individual-segregant datasets from SC medium (Table 1). Of these, three were mRNA datasets [13,15,45] and four were protein datasets [45,49,55,59]. To ensure balance between mRNA and protein, genes were included if they were present in at least two mRNA datasets and two protein datasets, for a total of 385 genes.

We conducted two genome-wide scans (Supplementary Table S6). The first scan searched for “joint QTLs” with additive signal across datasets. This scan revealed 868 joint QTLs at an FDR of 5%, with effects on the expression of 381 unique genes. Genes had a range from zero to five joint QTLs, with a median of two (Table 2).

**Table 2.** QTLs detected in joint mapping.

| QTL type | joint | consistent | Suggestive interaction | Strict interaction |
| --- | --- | --- | --- | --- |
| Number | 868 | 719 | 149 | 64 |
| Number of genes (out 385 tested) | 381 | 350 | 131 | 64 |
| Range per gene | 0 – 5 | 0 – 5 | 0 – 3 | 0 – 1 |
| Median per gene | 2 | 2 | 0 | 0 |
| Local | 48 | 32 | 16 | 5 |
| *Trans* | 820 | 687 | 133 | 59 |

Joint QTLs can reflect consistent effects that are similar across datasets, or a strong effect in some datasets with no or weak effect in the other datasets. To focus on joint QTLs with consistent effects on both gene products, we conducted *post hoc* interaction tests at the peak markers of joint QTLs. A joint QTL was classified as “consistent” if it showed no evidence for different effects on mRNA and protein (interaction test between genotype and gene product, nominal p-value > 0.05). In contrast, “suggestive interaction QTLs” showed tentative evidence for different effects on mRNA versus protein at a nominal interaction p-value < 0.05. Of the joint QTLs, 719 were consistent QTLs (Table 2). About 4% of the consistent QTLs (n = 32) were local, with the remaining 96% acting in *trans*.

To further examine how the consistent QTLs affected mRNA and protein, we split the datasets by gene product and separately tested for effects at the QTL peak marker. This showed that 79% of the consistent QTLs had significant effects (likelihood ratio test (LRT), nominal p-value < 0.05) on both mRNA and protein with the same direction of effect (Table 3). Some consistent QTLs showed significant effects only on mRNA (n = 62, 9%) or only on protein (n = 74, 10%), suggesting incomplete power of either the *post hoc* interaction test and/or of the LRT for the non-significant gene product. A small number of consistent QTLs (n = 14, 2%) were significant for neither mRNA nor protein, suggesting they reflect weaker effects that were only detectable due to the increased statistical power from combining multiple datasets.

**Table 3.** Classes of directional agreement of QTLs detected in joint mapping.

| mRNA significant<sup>1</sup> | Protein significant<sup>1</sup> | Same direction | Consistent QTLs (n = 719) | Strict Interaction QTLs (n = 64) | Interpretation |
| --- | --- | --- | --- | --- | --- |
| yes | yes | yes | 569 (79.1%) | 4 (6.3%) | Same Direction |
| yes | yes | no | 0 | 37 (57.8%) | Opposite direction |
| yes | no | NA | 62 (8.6%) | 9 (14.1%) | RNA-specific |
| no | yes | NA | 74 (10.3%) | 14 (21.9%) | Protein-specific |
| no | no | NA | 14 (1.9%) | 0 | Not detectable in single studies |

<sup>1</sup>based on LRTs at the peak marker performed separately in datasets of the given gene product

The second scan directly searched for loci with different effects on mRNA and protein, indicated by genome-wide significant interaction terms (“strict interaction QTLs”). In particular, this test can detect loci at which the effects on mRNA and protein have opposite directions. In this arrangement, the effects may cancel out in the first scan above, precluding discovery. The second scan revealed 64 strict interaction QTLs at 5% FDR, one for each of 64 genes (Table 2). The fraction of strict interaction QTLs that were local (n = 5, 8%) was not statistically different from that in the consistent QTLs (Fisher’s exact test (FET), p = 0.2). The strong majority of strict interaction QTLs (92%) acted in *trans*.

Most of the strict interaction QTLs (n = 37, 58%) had significant effects on both mRNA and protein, but with opposite direction of effect (Table 3). None of the consistent QTLs showed this pattern, as expected. More strict interaction QTLs were protein-specific (n = 14, 22%) than mRNA-specific (n = 9, 14%); each of these two categories was slightly more frequent among the strict interaction QTLs than among the consistent QTLs. Four strict interaction QTLs had significant effects on mRNA and protein in the same direction. Here, the interaction effect reflects a quantitative difference in how strongly the locus affects mRNA vs protein.

The strict interaction QTLs were enriched at several hotspot locations (Figure 5A). Most occurred at the *MKT1* locus (36 QTLs, 56%). This region also included 70% (26 / 37) of those strict interaction QTLs with opposite direction of effect on the two gene products, with the remainder distributed across the genome. Several strict interaction QTLs occurred in a region on chromosome XIII (11 QTLs, 17%) that contains *BUL2*, where a missense variant between BY and RM causes a difference in amino acid uptake [60].

We conducted a separate search for QTLs that only affected mRNA or only affected protein by comparing the QTLs identified by individual mapping in all 13 datasets, including all growth media and the bulk-segregant QTLs (Supplementary Text 3; Supplementary Table S7). The resulting mRNA-specific or protein-specific loci were also concentrated at *MKT1* and *BUL2*, in addition to *IRA2* and *LEU2* (Figure 5A).

We used mashr posterior summaries (Methods; Supplementary Table S8) to further examine the classes of directional agreement among QTLs identified in our two joint mapping scans. Among the 631 joint and strict interaction QTLs that had the same direction of effect on mRNA and protein using LRTs (Table 2), 579 (94.9%) also had concordant sign in mashr (see Figure 5B for an example). On the other hand, only 11 of the 37 (29.7%) strict interaction QTLs with opposite sign were recapitulated as discordant by mashr, a significantly lower rate of agreement between mashr and standard joint mapping (Fisher’s exact test p-value < 2.2e-16). Figure 5C-E shows examples of strict interaction QTLs with opposite effects or gene product-specific effects. There was no strict interaction QTL at which mashr detected effects that were significant (lsfr < 0.05) and in opposite directions across all datasets. These results indicate that directional agreement of genetic effects on mRNA and protein is a predominant and robust signal, whereas discordant effects are fewer and less robustly detected.

Together, these results revealed QTLs with significantly different effects on mRNA vs. protein of individual genes. The number of such QTLs is more than an order of magnitude lower than that of consistent QTLs with similar effects on the two gene products. We acknowledge that the additional statistical power needed to detect interaction QTLs may have reduced their number. The majority of strict interaction QTLs had significant effects on both mRNA and protein, but with opposite direction of effect. QTLs that exclusively affected only protein or only mRNA, with no effect on the other gene product, were a minority. Crucially, the *trans*-acting hotspots at which interaction QTLs occurred also influenced other genes in a manner that was *not* specific to mRNA or protein. Thus, *trans*-acting hotspots have varied effects on mRNA and/or protein levels of individual genes, with more gene product-specific effects at some hotspots than others.

## Discussion

To examine similarity between genetic effects on mRNA abundance vs. protein levels, we conducted an integrative analysis of 13 published datasets generated in a cross of the BY and RM yeast strains. Multiple observations indicated high concordance between the genetics of mRNA and protein. First, global sharing of genetic effects was high across all datasets, with no obvious separation of mRNA and protein datasets. Second, the genetic architectures of mRNA and protein levels were similar in that both were shaped by similar numbers of local QTLs and *trans*-acting QTLs. Third, the level of agreement between eQTLs that shape mRNA abundance and pQTLs that shape protein levels was no different from the level of agreement among eQTL or among pQTL datasets. Fourth, *trans*-eQTLs and *trans*-pQTLs were clustered at the same hotspot positions, and no hotspot was purely specific to mRNA or protein. Fifth, joint genetic mapping across datasets revealed an order of magnitude more QTLs with consistent effects on mRNA and protein than QTLs with significantly discordant effects. Together, these results suggest that regulatory variation typically has the same effects on mRNA and protein.

Against this backdrop, we did identify loci whose effects on mRNA and protein of the same gene appear to differ. Specifically, *trans*-acting hotspots at the *MKT1* and *BUL2* genes were enriched for jointly-mapped interaction QTLs, with particularly strong enrichment of QTLs with opposite sign on mRNA and protein at *MKT1*. In a separate analysis, *MKT1* and *IRA2* were also enriched for mRNA-specific effects, while *BUL2* was enriched for protein-specific effects. Our results broadly agree with recent work [5,21,37,45] suggesting that many genetic effects on mRNA and protein are consistent, while some *trans*-acting hotspots tend to have more gene product-specific effects than others. Future work will need to examine the precise quantitative mechanisms through which these hotspots affect the transcriptome and the proteome in ways that yield consistent *trans* effects on the mRNA and protein for many genes but also effects on just mRNA, just protein, or discordant effects for other genes.

Our reanalysis has implications for future studies into the two layers of gene expression. We showed that comparing a dataset of a given gene product to just one dataset of the other gene product can be misleading. Depending on which dataset is chosen for comparison, the results might appear to show substantial gene product specificity, even though comparison to another dataset might have shown more agreement. Two datasets of the *same* gene product can differ considerably (Figure 3), probably due to a combination of incomplete statistical power, uncontrolled environmental variation, and other unavoidable idiosyncrasies of any given study. Indeed, when exploring these data, we noticed many QTLs that would have appeared to be specific to one gene product in a comparison between just two datasets but where consideration of additional datasets then revealed the “missing” QTLs in the other gene product. The most parsimonious explanation for this situation is that these QTLs do in fact have concordant effects on both gene products. Ideally, future datasets would be compared to multiple existing datasets to gauge the reproducibility of any potential discrepancies. We recognize that the BY/RM cross is unusual in the large number of regulatory variation studies that are available. In other systems with less available data, strong conclusions of discordant effects might require experimental verification, for example by dissection of causal variants followed by measuring their effects on the two layers of gene expression.

Context-specificity and gene-by-environment interactions (GxE) are known to be important for regulatory variation [23,26]. When mRNA and protein are measured in non-identical conditions, environmental interactions can create the impression of gene product-specificity. The wide variation we observed among studies of the same gene product, even among those purposefully performed in the same medium, suggests that GxE due to subtle, unintentional environmental differences may be hard to control in separate studies. To avoid any opportunity for environmental confounding, studies of mRNA and protein should seek perfect environmental control by using the exact same growth cultures for both mRNA and protein. Two studies have recently implemented this rigorous design [45,51], but remained limited in the number of segregants or the number of genes they could analyze.

Our analyses have several limitations. First, low statistical power is a concern for nearly all datasets we examined. Many QTLs go unnoticed in underpowered studies, reflected in the fact that more highly powered studies have revealed many previously “missing” QTLs that increased the observed concordance between the genetics of mRNA and protein [5,37]. Second, heterogeneity in the environment used in each study may have influenced our results. For example, because there is no individual-segregant protein dataset in YNB, joint QTL mapping could not be performed in a fully crossed fashion, with both mRNA and protein measured in both SC and YNB. The BY/RM cross still lacks a study in which many genes are measured for mRNA and protein expression in the same experimental cultures and in a large enough sample to account for most genetic variation. Third, as in any genetic mapping design based on meiotic recombination, linkage among neighboring variants may have influenced our results by blending the effects of any linked causal DNA variants. Finally, yeast is not well suited for modeling the effects of protein secretion and protein-specific extracellular processes such as modification or uptake by organs that influence human blood or plasma proteomes [30]. Instead, yeast is an excellent model for intracellular dynamics of gene expression processes.

In summary, we have presented evidence consistent with a parsimonious null model under which genetic effects on mRNA and protein are largely concordant. Several loci with discrepant effects are notable exceptions to this global trend and warrant further studies. Our results highlight crucial challenges for future work on the effects of regulatory variation on mRNA vs. protein. Going forward, we emphasize the importance of perfect environmental control through measuring mRNA and protein in the same cultures, of high statistical power to avoid false impressions of gene product-specificity through missed QTLs, and of experimental dissection to verify and understand the basis of any effects that are specific to mRNA or protein. Based on the results here, we expect that any discrepancies in the effects of regulatory variation on mRNA and protein will be subtle and quantitative rather than strongly specific to mRNA or protein.

## Materials & Methods

Unless otherwise noted, all analyses were performed using the R programming language version 4.5.2 [61]. Various analyses used the tidyverse, ggplot2, RColorBrewer, and pheatmap packages [62–65]. Analysis code is available at https://github.com/Krivand/BYxRM_jointMapping

### Data sources

Microarray data for the Brem et al., 2002 [13] and 2005 [15] studies were retrieved from the NCBI GEO curated dataset browser (https://www.ncbi.nlm.nih.gov/sites/GDSbrowser/). The accession numbers for the Brem et al., 2002 datasets are GDS91 and GDS92. The accession numbers for the Brem et al., 2005 dataset are GDS1115 and GDS1116. mRNA abundance data from Smith & Kruglyak 2008 was obtained from Dataset S2 in [23]. Protein quantifications from Khan et al., 2009 [59] were kindly provided by Joshua Bloom and are available here as Supplementary Table S9. Protein abundances from Foss et al., 2011 were from Table S1 in [49]. Data from Picotti et al., 2013 were from the “SRM-dataset” tab in the Supplementary Data file from [55]. Genotype and covariate data from Grossbach et al., 2022 were obtained from Dataset EV2, and mRNA and protein abundances were obtained from Dataset EV1 in [45]. For Albert & Bloom et al., 2018, gene expression, covariate, and genotype data were obtained from Source data 1, 2, and 3 in [5], respectively. Distant pQTLs from Albert et al., 2014 were obtained from Supplementary Data 2 in [37]. Local pQTLs from that paper were obtained from their Supplementary Table S2. QTLs from Brion et al., 2020 were obtained from Source Data 2 (pQTLs) and Source Data 3 (eQTLs) in [51].

### Genotype data processing

#### Segregant genotyping

Genotypes for the segregants from Albert & Bloom et al., 2018 were used as published [5]. The segregants used in Brem et al., 2002, Brem et al., 2005, Smith & Kruglyak 2008, Khan et al., 2009, Foss et al., 2011, and Picotti et al., 2013 had been re-genotyped by incorporation of RNA-sequencing data by Grossbach and colleagues [45]. Here, we matched the segregants in the earlier datasets with the more-recent genotypes provided by Grossbach et al.. We coded BY and RM alleles to have values of -1 and 1, respectively. Therefore, positive QTL effects reported here indicate higher expression from the RM compared to the BY allele.

#### Imputation of missing genotypes

In the Grossbach et al., 2022 data, some segregants had missing genotypes at a small number of the 3,593 markers (median of 1 missing marker per segregant; range: 0 – 12). We imputed these missing genotypes as follows. The imputed genotype was -1 or 1 if the upstream and downstream markers had the same genotype. The imputed genotype was 0 if the flanking markers had different genotypes. This intermediate genotype value reflects that a recombination event must have occurred at an unknown position within the stretch of missing genotypes. If a stretch of missing genotypes was located at the end of a chromosome, these markers were assigned the genotype of the nearest genotyped marker.

### Phenotype data

#### Missingness control and imputation

For all datasets, we removed any genes or segregants with more than 25% missing phenotypes in the given dataset. We imputed any remaining missing phenotypes with one of two methods. Imputation was needed because the QTL mapping framework used here requires complete data matrices. Imputation also maintained consistent sample sizes across genes in a given dataset.

##### Imputation method 1

For datasets without covariates (Brem et al., 2002, Brem et al., 2005, Smith & Kruglyak 2008, Khan et al., 2009, Foss et al., 2011, and Picotti et al., 2013), we imputed missing values with their gene-wise means across observed samples.

##### Method 2

For phenotype data from Grossbach et al., 2022, which had covariate information, missing values were imputed using a two step process at each gene with at least one segregant that lacked phenotype information. First, we built linear models of phenotypes as a function of covariates using all available phenotypes and covariates. Second, we imputed missing phenotypes for a given segregant by using these models to predict the missing expression value from the given segregant’s covariates. In cases where the effects of a level of a given covariate could not be estimated because the data points representing that level were all missing, we built the model without that covariate.

#### Dye swap channel averaging

From the dye swaps used in Brem et al., 2002 and 2005, the mean blank signal was subtracted, and the two swaps were averaged to yield a single log expression ratio per segregant relative to the reference strain. Sign was inverted so that values correspond to log(RM/BY).

#### Strain replicates, phenotype replicates, and covariate correction

The Picotti et al., 2013 dataset had two replicates for 5 proteins and 2 segregants. We processed these measures by first averaging them per protein and then per segregant. This process yielded measures from 48 proteins and 76 strains.

The Grossbach et al., 2022 datasets had replicated measurements for some segregants. Specifically, for the mRNA measurements, 83 strains had 1 replicate, 25 strains had 2 replicates and 2 strains had 3 replicates. For the protein measurements, 83 strains had 1 replicate, 25 strains had 2 replicates and 6 strains had 3 replicates. Grossbach et al., 2022 also provided covariates for both mRNA and protein measurements. To account for covariates while using all replicates, we took two different approaches for processing the data for individual versus joint QTL mapping.

##### Individual mapping

When the replicates of a segregant had different covariate values, we averaged their phenotypes to create an aggregated segregant. We then assigned covariate values to this aggregated segregant by computing weighted fractional covariate membership proportional to the covariates of the individual replicates. For example, if two replicates of a segregant were in a batch A and the third was in batch B, the aggregated segregant was assigned membership to batch A of 2/3, and membership to batch B of 1/3. We then used a linear model to perform covariate correction on all strains.

##### Joint mapping

The joint mapping pipeline cannot accommodate dataset-specific covariates. Therefore, we used a linear model to perform covariate correction on each replicate of every segregant. We then averaged all replicates from a given segregant to form one set of phenotypes for that segregant.

#### Removal of parental strains

If present, data from the BY and RM parent strains were removed from the genotype and phenotype matrices prior to further processing.

### QTL mapping for individual datasets

QTL mapping in individual datasets was performed as described previously [5]. Briefly, the approach uses a per-chromosome pre-correction and an iterative two-phase mapping. The per-chromosome pre-corrections improve the signal to noise ratio when mapping on a specific chromosome by removing strong individual genetic effects from other chromosomes as well as accounting for the overall genetic relatedness of the other chromosomes. When present, covariates were corrected out prior to this analysis, leaving residuals centered at zero. For datasets without covariates, phenotype values were regressed on a column vector of 1s to mean-center them. Phenotype values were then standardized gene-wise by subtracting the mean across segregants and dividing by the standard deviation. Genome scans were performed using these standardized phenotypes.

The power and precision of QTL mapping improves when genetic effects arising from other chromosomes are controlled for [5,66–68]. To control for genetic contributions from other chromosomes, we first performed a QTL scan in each individual dataset by calculating Pearson correlations between standardized genotypes and standardized phenotypes for each gene-marker pair (**eq. I**, see below), converting these correlations to LOD scores (**eq. ii**), and identifying up to three of the largest genetic effects from each chromosome with LOD > 3.5. These strong genetic effects from all other chromosomes are removed from the expression data on a per-chromosome basis by taking the residuals from a linear model that included all strong genetic effects from all the other chromosomes as fixed effects (**eq. iii**). For each chromosome, a genetic relatedness matrix was constructed using all markers from all other chromosomes, and the corresponding best linear unbiased predictors (BLUPs) of this polygenic background were estimated and subtracted. The resulting residuals were then used for QTL mapping on the focal chromosome (**eq. iv**).

Individual QTL mapping was performed in two iterated phases. In the scanning phase, gene-marker associations were calculated at all phenotype-marker combinations. In the permutation phase, 1000 permutations were performed in which LODs were calculated for phenotypes permuted relative to genotypes. FDR was calculated empirically by dividing the mean number of QTLs across genes discovered in each of the 1000 permutations by the number of QTLs discovered in the observed data. FDR was evaluated at LOD thresholds ranging from 1.5 to 9 in steps of 0.05. The LOD threshold corresponding to FDR = 5% was used to determine significant associations. Because permutation-based FDR estimates become unstable when very few peaks are present, scans were terminated when fewer than three peaks were detected.

Equations used in QTL mapping were as follows:

#### i. calculation of Pearson correlation at a gene-marker pair

Let *n* be the number of segregants, **Y** be the standardized phenotype matrix (segregants by gene expression phenotypes), and **X** be the standardized genotype matrix (segregants by markers). The matrix of pearson correlations **R**, where each element is the pearson correlation R between the phenotypes and genotypes across the segregants, is given by:

***R*** = ***Y****^T^**X**(n-1)^-1^*

#### ii. calculation of LOD score for a gene-marker pair

*LOD = -0.5n(log_10_(1-R^2^))*

Here, *R* is the Pearson correlation between the genotype and phenotype and *n* is the number of phenotypic observations.

#### iii. per chromosome removal of strong additive effects from other chromosomes

*E* ∼ *q_!i_* + *S_i_*

Here, *E* is the expression of a gene across segregants. Each chromosome *i* serves as the focal chromosome while searching for QTLs on it. For this search, marker genotypes at previously identified strong QTLs on all other chromosomes (*q_!i_*) are included as additive covariates. We then obtain the residuals *S_i_*, which are expression phenotypes corrected for *q_!i_*.

#### iv. per chromosome removal of polygenic additive effects from other chromosomes

*S_i_* ∼ *p_!i_* + *T_i_*

Here, *p_!I_* is the polygenic effect from a genetic relatedness matrix constructed using all markers from all other chromosomes, estimated as BLUPs. *T_i_* are the final residuals after removal of both strong additive effects (via **eq. iii**) and the polygenic background. These residuals are used for QTL mapping on chromosome *i*.

### QTL confidence intervals

Confidence intervals were defined as the contiguous marker interval surrounding the peak marker with LOD scores no more than 1.5 lower than the peak LOD value. Markers reported in [45] correspond to stretches of BY/RM variants in perfect linkage disequilibrium. Therefore, for QTLs mapped in datasets using these genotypes, we extended confidence intervals to include BY/RM those variants in the given stretch that are furthest from the QTL peak.

### Classification of QTLs as local vs *trans*

QTLs were classified as local if the location of the gene affected by a QTL overlapped the QTL by at least one nucleotide. For this classification, QTL confidence intervals were extended by 10 kb. This was done to avoid incorrect *trans* classifications for very strong local QTLs, some of which are so narrow that they exclude the gene’s open reading frame even though they reside just upstream or just downstream of their target gene, suggesting *cis*-regulatory effects on the promoter or 3’UTR. Gene locations were based on Ensembl [69] version 83, which corresponds to the sacCer3 *Saccharomyces cerevisiae* genome build [70].

### Comparison to published QTLs

#### Sources and preprocessing of published QTL lists

Published QTLs were available from the following publications:

For Smith *et al.* 2008, QTLs were downloaded from Table S4 in [23]. Published QTL coordinates were transferred to sacCer3 using the UCSC Genome Browser liftover tool [71]. Effect sizes were calculated by subtracting the reported “RM effect” from the “BY effect” in the given medium.

For Foss *et al.* 2011, QTLs were downloaded from Table S4 in [49]. QTL peak marker positions were transferred to sacCer3 using liftover. Intervals of QTL positions were not available. Instead, we used the median QTL width observed in our re-mapping of these data (51,314 bp) and centered intervals of this size on the published peak markers. Effect sizes were not available for these published QTLs.

For Grossbach *et al.*, 2022, QTLs were downloaded from Dataset EV4 in [45]. Effect sizes were not available. For Albert *et al.* 2018, QTLs were downloaded from Source Data 4 in [5].

#### Comparison between remapped and published QTLs

We used the R GenomicRanges package [72] to assess whether a given remapped QTL overlapped a published QTL for the same gene, and *vice versa*. Where available, we required the same sign of effect in both QTL sets. We conducted this analysis for all QTLs as well as the 20% strongest QTLs, as ranked by LOD score. For overlapping QTLs, we computed Spearman rank correlations of effect sizes, where available. For these correlations, QTLs were deemed to overlap irrespective of their sign.

### QTL discovery rates as function of sample size, gene product, and mode of action

Statistical analyses in this section were restricted to individual-segregant datasets, excluding bulk segregant datasets. To explore patterns in QTL discovery, we fit the numbers of mapped QTLs across studies using a negative binomial linear model as implemented via the glm.nb() function in the MASS package [73]. Sample size was included as a predictor after transformation to log_10_. The logarithm of the number of genes analyzed in each dataset was included as an offset. Linear models were examined using analysis of variance (ANOVA) to compute p-values. For analyses comparing rates of discovery for local vs *trans* QTLs, we used the glmmTMB package [74] to fit a mixed negative binomial linear model that incorporated dataset identity as a random term to ensure the numbers of local and *trans* QTLs from a given dataset were properly paired.

### Hotspot analyses

Locations of QTLs were visualized in Figure 4 using the hist() function in R. To examine how many mRNAs or proteins a given hotspot affected, we divided the genome into non-overlapping bins of width 100 kb for visualization in Figure 5 or 10 kb for more fine-grained analyses (Supplementary Table S5). The number of QTLs from a given analysis whose peak markers were located in a given bin were counted. We expressed the strength of a given bin as the fraction of QTLs from that dataset that occurred in the bin.

### Pairwise comparisons between datasets

QTL overlap comparisons were performed using the GenomicRanges package [72]. QTLs were padded by 10 kb to avoid missing overlaps between strong QTLs, whose small confidence intervals may otherwise narrowly miss each other. These pairwise comparisons between datasets are not symmetric. Therefore, a single pair of datasets yields two observations, with each dataset serving once as the query and once as the target.

We analyzed only loci that can exist in both datasets in a given pair. For example, the mating type locus affects the expression of several genes in *trans*, but can only do so if a dataset has segregants of both mating types (Supplementary Table S1). Thus, the mating type locus was analyzed only when comparing studies with both mating types present. In addition to the mating type, loci excluded from some dataset pairs were *LEU2*, *URA3*, *AMN1*, and *KRE33*.

The bulk segregant datasets were only designed to detect local effects for genes at which both parent strains had been engineered with the respective fluorescent markers. Therefore, all local QTLs were excluded from analyses with Brion et al., 2020 [51] as the target dataset. When the bulk-segregant dataset from Albert et al., 2014 [37] was the target dataset, local QTLs were only included for the 41 genes that had been tagged in both parents in [37].

Distributions of overlap fractions were compared using mixed binomial linear models fit using the glmer function from the lme4 package [75]. QTLs from a query dataset with a match in the target dataset were treated as hits, and those without a match were treated as misses. Hit fractions were modelled as a function of gene product pair (i.e. mRNA-to-protein, protein-to-mRNA, mRNA-to-mRNA, or protein-to-protein) as fixed effects, with separate models comparing one gene product pair to one other gene product pair (e.g. mRNA-to-protein compared to protein-to-mRNA). The models included random terms controlling for the identity of the query and target datasets. This was necessary because there were systematic differences in overlap fractions among datasets. For example, using the large Albert & Bloom et al., (2018) dataset [5] as a query tended to yield low agreement with all other datasets as targets, irrespective of whether these targets were for mRNA or protein. This is because the higher power of this dataset enabled identification of many weak eQTLs that cannot be detected in other studies due to statistical reasons.

To compute the π1 statistic, we took the peak marker of a given QTL in the query dataset and extracted the raw p-value from the QTL mapping scan at the same marker in the target dataset. Target dataset p-values were from the initial scan for that dataset, without correction for QTLs on other chromosomes. We then computed π1 on the resulting p-value distribution using qvalue [57]. Bulk-segregant datasets were excluded as target (but not as query) datasets for π1 analyses because p-values at individual markers are too noisy in these datasets, which rely on average signals across neighboring markers. Only query datasets with at least 80 QTLs were analyzed for π1 to ensure sufficient numbers of p-values to fit the underlying model. In rare cases, the p-value distribution lacked values close to 1, which prevented the π1 algorithm from running. In these cases, we appended one p-value equal to 1 to the p-value distribution. The π1 statistics for different gene product pairs were compared using mixed linear models with query and target datasets as random variables.

### QTLs with specific effects on mRNA or protein of specific genes

QTLs overlap was assessed using the GenomicRanges package [72].

### QTL joint mapping

#### Porting Albert & Bloom et al. genotypes into the Grossbach et al. marker coordinate system

To enable joint mapping analyses, we ported the genotypes of the segregants in Albert & Bloom et al., 2018 into the Grossbach et al. 2022 marker set used for QTL mapping in all the other individual-segregant datasets. Each segregant in Albert & Bloom was assigned a genotype for a given Grossbach marker, using the value of the nearest Albert & Bloom marker. When multiple Albert & Bloom markers mapped to the same Grossbach marker, their genotype values were averaged, sometimes yielding fractional values between −1 and +1. This procedure resulted in a genotype matrix representing all segregants in Grossbach et al. 2022 marker space for joint QTL analyses.

#### Inclusion criteria for joint QTL mapping

To ensure that genotype-by-gene product interaction effects were estimable and not driven by single-dataset outliers, we required that each tested gene be measured in at least two mRNA and two protein datasets. Using these criteria, 385 genes were eligible for joint mapping.

#### Joint Mapping

Joint QTL mapping was performed across datasets by testing for marker associations using likelihood ratio tests in linear mixed models. For each gene, phenotype observations were used from all datasets in which that gene was measured. Within each dataset, phenotypes were standardized gene-wise to mean zero and unit variance prior to joint analysis. These within-dataset standardized values were then concatenated across datasets for a given gene to form the phenotype vector used in the joint models; no additional cross-dataset standardization was applied. Phenotypes were pre-corrected for strong background QTLs identified in the individual dataset scans described above, removing large additive effects from other chromosomes prior to joint association testing.

For each marker and gene, we fit two nested linear mixed models with dataset-specific intercepts and compared them by likelihood ratio tests (LRT). When no interaction effect was modeled (i.e., “joint QTL” discovery), the null and alternative models were:

*m0: : pheno ∼ (1 | dataset)*

*m1: : pheno ∼ (1 | dataset) + geno*

Here, *pheno* is a vector of expression values for the given gene for each segregant, *dataset* is a random variable that controls for any systematic differences between datasets, and *geno* is a vector of genotypes for each segregant at the given marker.

When an interaction effect was modeled (i.e., “strict interaction QTL” discovery), the genotype main effect was included in the base model and the interaction term was tested by comparing the following models:

*m0: pheno ∼ (1 | dataset) + analyte + geno*

*m1: pheno ∼ (1 | dataset) + analyte + geno + geno:analyte*

Here, “analyte” is either mRNA or protein.

For each gene-marker pair, the LRT p-value from comparing m1 to m0 was recorded, and a LOD score was computed from the log-likelihoods as:

*LOD = (logL(m1) − logL(m0)) / log(10)*

For joint QTLs, the reported effect size was the fitted genotype coefficient. For strict interaction QTLs, the reported effect size was the genotype-by-analyte interaction coefficient.

To enable detection of multiple independent QTLs per gene, mapping was performed in iterative passes. After each pass, the most recently discovered QTL for each gene was regressed out by refitting the corresponding alternative model (*m1*) at the peak marker and replacing the gene’s phenotype with the residuals from this model. Residualized phenotypes were then re-tested in the next pass using the same procedure. Passes continued until no significant associations remained (see below).

For each discovered association, we defined a marker interval on the chromosome using a 1.5 LOD-drop criterion as described for QTL mapping from single datasets.

#### FDR control

Because joint QTL mapping required fitting large numbers of linear mixed models, and because each gene was supported by a different combination of contributing studies, genes were processed in chunks for tractability.

In each chunk, genes were processed in passes of an iterative forward-search for QTLs. Within each pass, p-values across all tested gene-marker pairs in the chunk were adjusted using the Benjamini-Hochberg (BH) procedure [76]. In this implementation, each pass defines a separate set of hypotheses for FDR control within that chunk. Because BH adjustment depends on both the number of tested hypotheses and the rank ordering of p-values within the tested set, applying BH separately within chunks yields adjusted p-values that differ slightly from those obtained under a single global correction across all genes. Chunk sizes were kept similar to reduce sensitivity of BH-adjusted p-values to the total number of hypotheses in each correction. Associations with BH-adjusted p-values less than or equal to a threshold of 0.05 were considered significant. Within each pass, we selected the marker with the minimum p-value among BH-significant hits, separately for each gene.

Chunk sizes were chosen to balance computational efficiency and stability of the BH adjustment. Specifically, genes were processed in 8 chunks of size 50, with the final chunk containing 35 genes.

#### Effects of jointly-mapped QTLs on each gene product

LRTs were performed as above but in only the mRNA or protein datasets, using these nested models:

*m0: pheno ∼ (1 | dataset)*

*m1: pheno ∼ (1 | dataset) + geno*

Because these analyses are descriptive and were performed *post-hoc* at QTLs discovered from genome-wide searches with rigorous FDR control, p-values were not corrected for multiple testing.

### Multivariate Adaptive Shrinkage (mashr)

#### Model fitting

We assembled matrices of effect estimates and corresponding standard errors across datasets, with rows representing gene–marker pairs and columns representing datasets. To ensure robust covariance estimation, we restricted training to the 385 genes present in at least two individual-segregant mRNA and two individual-segregant protein datasets also used in joint-QTL mapping above. Because the Albert & Bloom et al., 2018 dataset contained substantially more segregants than the other datasets, we calculated its association statistics in data downsampled to 100 segregants to prevent this single dataset from disproportionately influencing the learned covariance structure. To reduce redundancy from linkage disequilibrium, markers were LD-pruned based on Grossbach et al., 2022 genotypes [45] using an absolute correlation threshold of 0.8, leaving 433 markers. For model training, we used two distinct subsets of the 166,705 gene–marker pairs. Covariance matrices were estimated from 20,000 rows sampled with enrichment for strong signals (40% drawn from the top 20% by maximum absolute Z-score, with the remainder drawn from the background). Mixture weights were then estimated using all LD-pruned gene–marker pairs.

#### Quantifying global similarity

To quantify global similarity of genetic effects across datasets, we used the covariance structure learned by mashr [54]. Mashr represents effect sizes across studies as a mixture of covariance matrices that each describe a distinct pattern of effect sharing (e.g. effects present in all studies, only in one dataset, etc), with corresponding mixture weights (π). We constructed a single aggregate covariance matrix as the π-weighted sum of these covariance components, such that for each pair of studies (i, j), the entry in this matrix is the weighted average of their covariance across all sharing patterns, with weights given by π. This matrix represents the expected covariance between studies for a randomly selected genetic effect under the fitted mashr model. We then converted this covariance matrix to a correlation matrix, which normalizes for differences in effect size scale across datasets. The resulting matrix provides, for each pair of datasets, the expected correlation of genetic effects under the fitted mashr model. This correlation matrix is shown as a heatmap in Figure 1, where each entry reflects the similarity in effect direction and relative magnitude of genetic effects between two datasets, averaged genome-wide through the mashr mixture model.

#### Model evaluation

To test whether mashr assigns high-confidence effects to QTLs identified by joint mapping, we compared these QTLs to loci that were not identified as QTLs (here, “!QTLs”). !QTLs were defined as the set of markers not in linkage with any QTL peak marker at r > 0.8. The QTL and !QTL sets contained 453 and 1,139 markers, respectively. In each of 1,000 iterations, QTL and !QTL markers were LD-pruned separately using a random-order greedy procedure, in which markers were visited in random order, retained if not previously linked to a retained marker at r ≥ 0.8, and used to exclude all remaining linked markers. After LD pruning, the median number of retained markers across resampling runs was 165 for QTLs and 160 for !QTLs. For the gene–marker analysis, all gene–marker pairs involving retained QTL markers were included, whereas for each retained !QTL marker a single eligible gene–marker pair was randomly sampled. In both the QTL and !QTL sets we then calculated the fraction of gene–marker pairs with lfsr < 0.05, and this resampling analysis was repeated 1,000 times. At the gene–marker level, QTLs showed strong enrichment for lfsr < 0.05 compared to !QTLs (median across 1,000 iterations: 87.4% vs 9.4%). We performed an analogous analysis at the marker level by calculating the percent of markers for which at least one gene–marker pair had lfsr < 0.05, and observed a similar enrichment (89% vs 9.4%). These results indicate that jointly mapped QTLs are enriched for markers where mashr detects shared genetic effects across studies.

#### Model application to joint QTLs

To assess how QTLs identified in joint mapping behave under mashr, the fitted mashr model was applied to the non-LD-pruned peak markers identified in joint QTL mapping (453 unique markers) to compute posterior means of effect size and lfsr for each dataset. We then summarized their lfsr and effect direction across studies. Specifically, loci were classified as either “concordant” (each gene product showed at least one significant effect with lfsr < 0.05, all significant effects within each gene product had the same sign, and that sign was the same between gene product), “discordant” (each gene product showed at least one significant effect, all significant effects within each gene product had the same sign, but the signs were opposite between gene product), “specific” (one gene product had a significant effect in at least one dataset and the other had none), “mixed” (at least one gene product had significant effects in both directions), or “undetected” (all lsfr > 0.05). These mashr-derived directional classifications were compared to the corresponding “consistent” and strict interaction categories defined by joint QTL mapping (see Results for details on these categories) to evaluate agreement between the two approaches.

## Acknowledgements

We thank Joshua Bloom for providing the Khan 2009 protein quantifications. We thank members of the Albert & Gusev laboratories for comments on the manuscript. This work was funded by grant R35GM124676 from the NIH to FWA.

