## Supplementary Text, Figures, Table S4 for "High concordance between genetic effects on mRNA and protein abundance"

### Supplementary Materials – Table of Contents

**Supplementary Materials – Table of Contents** 1
**Supplementary Text** 2
Supplementary Text 1 2
Supplementary Text 2 2
Supplementary Text 3 3
Supplementary Text References 4
**Supplementary Figures** 5
**Supplementary Tables** 9

### Supplementary Text

#### Supplementary Text 1

##### QTL discovery in bulk-segregant versus individual-segregant datasets

We asked whether patterns of QTL discovery from individual-segregant panels were reflected in bulk-segregant studies. The median number of pQTLs per gene discovered in the two bulk segregant protein datasets by Albert et al. 2014 and Brion et al., 2020 [1,2] was not significantly different from eQTLs in the large Albert & Bloom individual-segregant dataset [3] (Figure 2A; paired Wilcoxon test of *trans*-QTL numbers for genes present in both datasets: $p \geq 0.13$), suggesting that these GFP-based implementations of the bulk segregant approach were about as statistically powerful as a panel of ~1,000 individual segregants. Bulk segregant mapping of RNA levels using a fluorescent reporter in Brion et al. [2] resulted in fewer QTLs than the 1,000 individual segregants from Albert & Bloom and the two protein bulk segregant studies ($p \leq 0.02$). While this lower discovery rate could indicate lower sensitivity of the fluorescent reporter compared to other mRNA quantification techniques, it may also reflect the fact that the reporter primarily measures RNA production rather than steady-state abundance of mRNA [2]. Genetic effects on mRNA stability and degradation [4] that create QTLs in typical eQTL studies would not be detectable using this reporter.

Of 41 proteins from the Albert et al. [1] bulk-segregant study for which local pQTL mapping had been conducted, 20 had a local pQTL. This is a similar fraction (49%) as that seen for local eQTLs in the large Albert & Bloom individual-segregant study [3] (Figure 2B). The same 41 proteins were affected by 347 *trans*-pQTLs in the bulk-segregant study, with a median of eight *trans*-pQTLs per gene. This was not significantly different from the median of seven *trans*-eQTLs observed for these genes in Albert & Bloom [3] (paired Wilcoxon test $p = 0.49$). Thus, both individual-segregant and bulk-segregant QTL mapping show prominent roles for *trans*-acting variation in both mRNA and protein abundance.

#### Supplementary Text 2

##### QTL agreement between and among mRNA and protein studies

Using pairwise agreement between QTLs of a given study pair, we found that comparisons between mRNA and protein datasets were not different from those among mRNA studies and among protein studies (Figure 3). We conducted a series of analyses to probe the robustness of this result (Supplementary Figure S3 and Supplementary Table S4).

To avoid potential issues with low statistical power to rediscover a given QTL in the target datasets, we first restricted the analyses to the 20% strongest QTLs in a given query set. As expected, agreements rose compared to using all query QTLs (Supplementary Figure S3 A &

B). However, the distributions of mRNA-to-protein and protein-to-mRNA comparisons remained statistically indistinguishable and did not differ from within-gene product comparisons ($p \geq 0.21$).

To avoid potential environmental confounding, we repeated the analyses using only pairs of studies conducted in the same growth medium. There were no statistically significant differences among the four gene product comparisons ($p \geq 0.21$). The same result was obtained when excluding comparisons involving bulk-segregant datasets ($p \geq 0.18$). Restricting the analyses to local QTLs resulted in overall stronger agreement between datasets (Supplementary Figure S3 C & D), but there was again no difference in replication rates between the four groups ($p \geq 0.18$).

Next, we estimated the fraction of query QTLs that replicated in a given target dataset by computing the $\pi 1$ statistic, using only individual segregant datasets as target sets (Supplementary Figure S3 E & F). Mean replication rates ranged from 42% (protein-to-protein) to 50% (protein-to-mRNA). Replication rates among the four comparison groups were not statistically different (mixed linear models: $p \geq 0.17$).

Finally, we computed correlations of effect sizes between query and target. Specifically, we computed Spearman's rank correlations between 1) the correlation coefficient between gene expression and genotype at each QTL peak marker in the query set with 2) the correlation coefficient between the same gene and the same marker in the target set, irrespective of the significance of association in the target set. We used the same set of markers as in the $\pi 1$ analysis. All of the 84 tested correlations were positive with a median of rho = 0.56, and all but two were at least nominally significant ($p < 0.05$). Restricting these analyses to the 20% strongest query QTLs typically resulted in stronger effect size correlations (median = 0.72, paired Wilcoxon test p-value = 1.0e-10). No comparison among the four groups was significant ($p \geq 0.10$; Supplementary Figure S3 G & H).

### Supplementary Text 3

#### Loci with consistent protein-specific or mRNA-specific effects on individual genes

Our joint QTL mapping was deliberately restricted to a subset of datasets due to two reasons. First, to avoid environmental confounding, we considered only datasets from the same growth medium. Second, bulk-segregant data is not compatible with the joint-QTL mapping pipelines.

In the analyses presented here, we loosened these restrictions to search for QTLs with effects that could be reproducibly specific to the mRNA or the protein of a given gene (Supplementary Table S7). We compared QTLs mapped individually in all 13 datasets for 451 genes that were present in at least two RNA and two protein datasets. Across all datasets, these genes were affected by 6,787 QTLs, with about half of these (3,559) from the large Albert & Bloom et al., 2018 eQTL dataset.

We searched for eQTLs that affected the mRNA of a given gene in at least two datasets with the same direction of effect and that had no pQTL for the same gene in any protein dataset. There were 307 such mRNA-specific eQTLs affecting 217 unique genes; 259 of these mRNA-specific loci (84%) were *trans*-acting. An analogous search for protein-specific pQTLs yielded 46 pQTLs affecting 38 genes. All protein-specific pQTLs were *trans*-acting.

We examined the locations of these *trans*-acting, gene product-specific QTLs (Figure 5A). The mRNA-specific QTLs were heavily enriched at the *MKT1* and *IRA2* hotspots. These hotspots do also affect numerous proteins, but for other genes. Their detection in the current analysis suggests that some of the effects of these hotspots on the mRNA of individual genes do not propagate to the protein level, adding further complexity to the wide-ranging effects of these loci.

The protein-specific pQTLs were enriched at different locations than the mRNA-specific eQTLs. The strongest enrichment (18 / 46 pQTLs) occurred at a region on chromosome III (70 - 140 kb), where the engineered *LEU2* auxotrophy segregates in some datasets and where strong *trans*-pQTL hotspots were seen in the Khan et al. and Foss et al. protein datasets [5,6]. We caution that this auxotrophy does not segregate in the large Albert & Bloom et al. eQTL dataset [3] and, thus, no eQTLs could be detected there. The Albert & Bloom et al. study did reveal “missing” eQTLs at many pQTLs that, without this large eQTL dataset, would have appeared to be protein-specific. If the *LEU2* auxotrophy had segregated in the large eQTL dataset, eQTLs may well have been discovered at this locus.

Another 16 protein-specific pQTLs occurred at 40 - 90 kb on chromosome XIII, which contains *BUL2*. The protein-specific pQTLs here were seen in a variety of protein datasets (rather than just two datasets for *LEU2*), and the *BUL2* missense variant does segregate in the large Albert & Bloom et al. eQTL dataset. While the *LEU2* and the *BUL2* loci both also affect the mRNAs of numerous genes (Figures 4 & 5), they do appear to have protein-specific effects on more genes than most hotspots.

#### Supplementary Text References

### Supplementary Figures

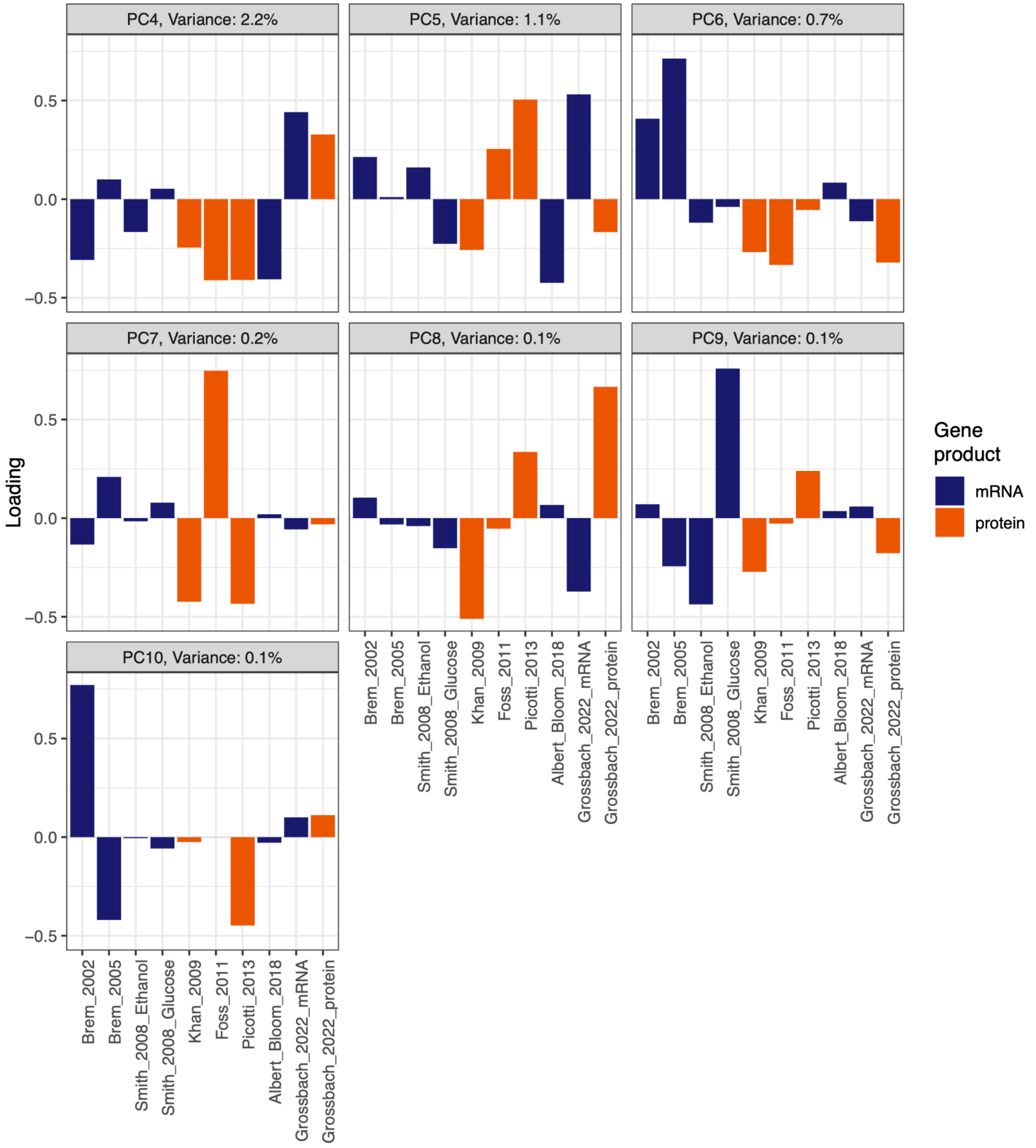


**Supplementary Figure S1.** Principal components (PCs) of mashr effect sharing. Loadings of each dataset onto each PC is shown on the y-axes. PC 4 – 10 are shown. PCs 1 – 3 are in Figure 1 in the main text.

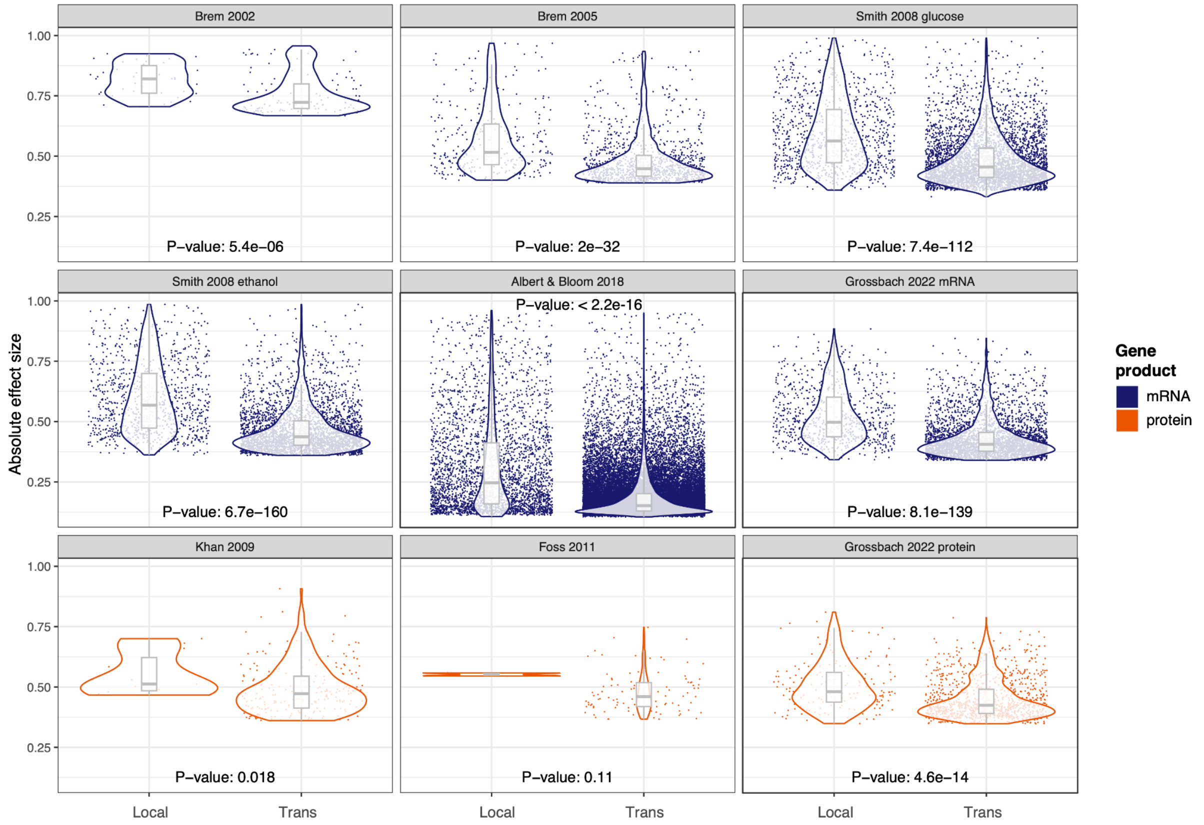


**Supplementary Figure S2.** Effect sizes of local and *trans*-QTLs. P-values for Wilcoxon rank tests comparing local and *trans* QTLs are indicated. P-values were not corrected for multiple comparisons. QTLs from Picotti *et al.*, 2013 are not shown due to their small number; the p-value for this dataset was p = 0.42.

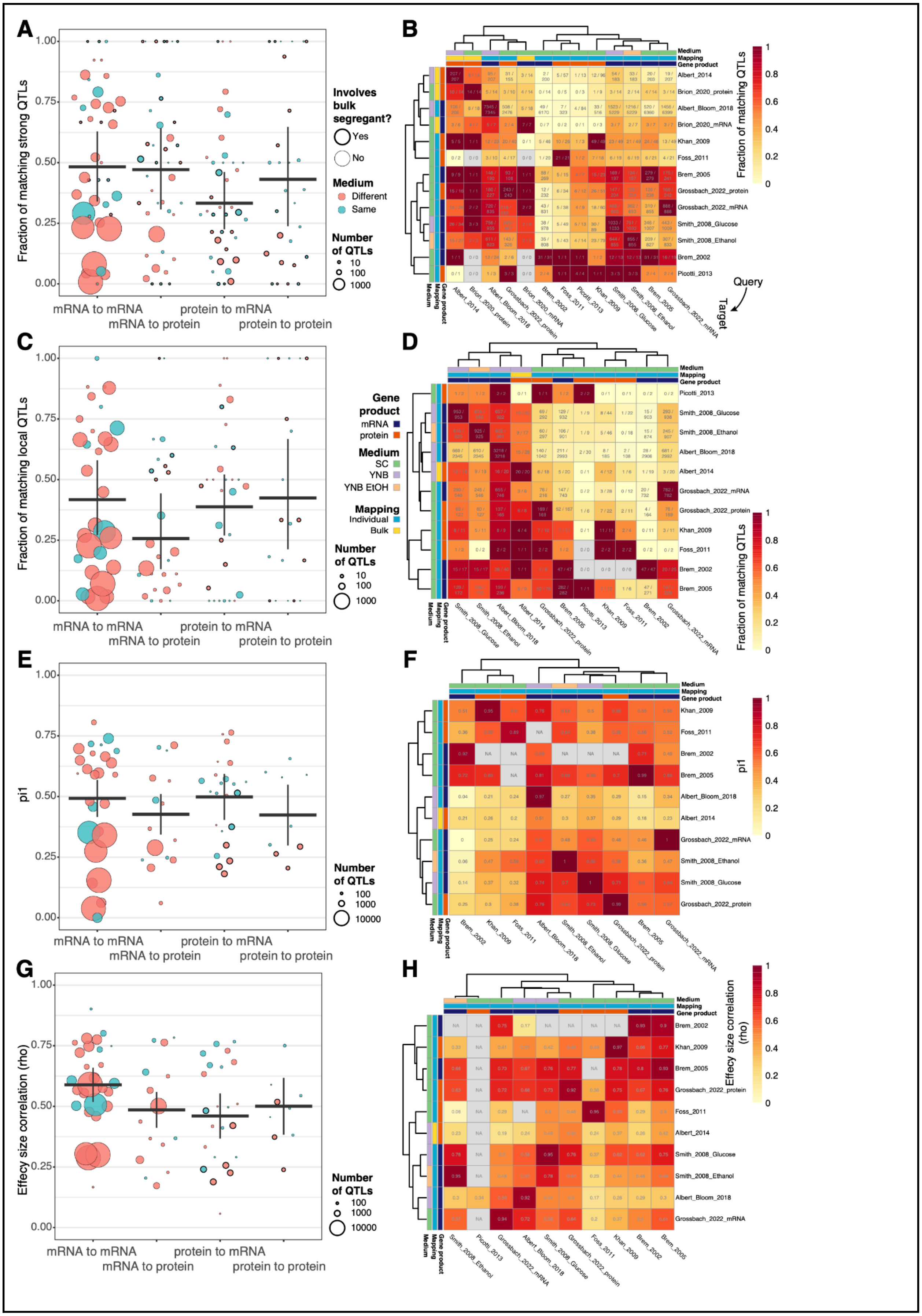
A
Fraction of matching strong QTLs
1.00
0.75
0.50
0.25
0.00
mRNA to mRNA
mRNA to protein
protein to mRNA
protein to protein
Involves bulk segregant?
Yes
No
Medium
Different
Same
Number of QTLs
10
100
1000
B
Medium
Mapping
Gene product
Fraction of matching QTLs
Albert_2014
Brion_2020_protein
Albert_Bloom_2018
Brion_2020_mRNA
Khan_2009
Foss_2011
Brem_2005
Grossbach_2022_protein
Grossbach_2022_mRNA
Smith_2008_Glucose
Smith_2008_Ethanol
Brem_2002
Picotti_2013
Query
Target
C
Fraction of matching local QTLs
Gene product
mRNA
protein
Medium
SC
YNB
YNB EtOH
Mapping
Individual
Bulk
D
E
pi1
100
1000
10000
F
G
Effecy size correlation (rho)
H

**Supplementary Figure S3.** Pairwise comparisons between datasets. A) Fraction of all QTLs in the query set that match a QTL in the target set, using only the 20% strongest QTLs in the query set. The figure shows the agreement for each pairwise comparison on the y-axis, with circles sized by the number of QTLs in the given comparison. Thicker circles denote a comparison involving a bulk-segregant study. Thick horizontal lines show the mean for each gene product comparison weighted by dataset sizes, as computed using linear models that control for dataset identity (Methods). Vertical lines are standard errors of these means, computed from the same models. B) The agreement for each pairwise comparison shown in A) is arranged as a heatmap. Each cell gives the number of query QTLs with a match in the target set and of all query QTLs. Gene product, mapping method, and growth medium are indicated; the color legend is in panel D. Rows show query datasets, and columns show target sets. Rows and columns are clustered based on similarity.
C) & D): As in A & B, but only for local QTLs. E) & F) As in A & B, but using the $\pi 1$ statistic as a measure of agreement. G) & H) As in A & B, but using the effect size correlations as a measure of agreement.

### Supplementary Tables

**Supplementary Table S4.** Comparisons between dataset agreements.

| Comparison[1] | All QTLs | Strong QTLs | Same medium | No bulk datasets | Local QTLs | π1 | Effect correlations |
|---|---|---|---|---|---|---|---|
| mm:mp | 0.97 | 0.69 | 0.51 | 0.52 | 0.14 | 0.42 | 0.10 |
| mm:pm | 0.53 | 0.21 | 0.30 | 0.94 | 0.76 | 0.45 | 0.26 |
| mm:pp | 0.51 | 1.00 | 0.93 | 0.63 | 0.89 | 0.87 | 0.13 |
| mp:pm | 0.57 | 0.59 | 0.21 | 0.74 | 0.45 | 0.17 | 0.85 |
| mp:pp | 0.53 | 0.90 | 0.52 | 0.74 | 0.18 | 0.86 | 0.93 |
| pm:pp | 0.41 | 0.39 | 0.48 | 0.18 | 0.92 | 0.48 | 0.72 |

The table shows p-values for contrasts between dataset agreements. [1]Each comparison contrasts the distribution of a given metric of agreement between datasets with the two indicated gene product pairings, where m: mRNA and p: protein. For example, “mm:mp” compares the distribution of agreement between two mRNA studies to the distribution of agreement between mRNA studies as queries and protein studies as targets. Note that agreements are not symmetric, such that mp ≠ pm. See text for further details.
